# *Mmp-13* deletion in cells of the mesenchymal lineage increases bone mass, decreases endocortical osteoclast number, and attenuates the cortical bone loss caused by estrogen deficiency in mice

**DOI:** 10.1101/2021.07.15.452452

**Authors:** Filipa Ponte, Ha-Neui Kim, Aaron Warren, Srividhya Iyer, Li Han, Erin Mannen, Horacio Gomez-Acevedo, Intawat Nookaew, Maria Almeida, Stavros C. Manolagas

## Abstract

The protective effect of estrogens against cortical bone loss is mediated via direct actions on mesenchymal lineage cells, but functional evidence for the precise molecular mechanism(s) and the mediators of these effects has only recently began to emerge. We report that the matrix metalloproteinase 13 (MMP-13) is the highest up-regulated gene in calvaria or bone marrow cells from mice lacking the estrogen receptor (ER) alpha in osteoprogenitors. We, therefore, generated mice with conditional Mmp-13 deletion in Prrx1 expressing cells (*Mmp-13*^ΔPrrx1^) and compared the effect of estrogen deficiency on their bone phenotype to that of control littermates (*Mmp-13*^f/f^). Femur and tibia length was decreased in sham-operated *Mmp-13*^ΔPrrx1^ mice as compared to *Mmp-1*3^f/f^. Cortical thickness and trabecular bone volume in the femur and tibia were increased and osteoclast number at the endocortical surfaces was decreased in the sham-operated female *Mmp-13*^ΔPrrx1^ mice; whereas bone formation rate was unaffected. Ovariectomy (OVX) caused a decrease of cortical thickness in the femur and tibia of *Mmp-13*^f/f^ control mice. This effect was attenuated in the *Mmp-13^ΔPrrx1^* mice; but the decrease of trabecular bone caused by OVX was not affected. These results reveal that mesenchymal cell–derived MMP-13 regulates osteoclast number, bone resorption, and bone mass. We have recently reported that the loss of cortical, but not trabecular bone, caused by OVX is also attenuated in *Cxcl12*^ΔPrrx1^ mice. Together with the present report, this functional genetic evidence provides proof of principle that increased production of mesenchymal cell-derived factors, such as CXCL12 and MMP-13, are important mediators of the adverse effect of estrogen deficiency on cortical, but not trabecular, bone. Therefore, the mechanisms responsible for the protective effect of estrogens on these two major bone compartments are different.

## INTRODUCTION

During the last 10 years, we and others have elucidated in genetic mouse models that the protective effects of estrogens on trabecular and cortical bone mass are mediated via ERα actions on distinct cell types: hematopoietic and mesenchymal lineage cells, respectively ^(1,2)^. In trabecular bone direct estrogen actions on osteoclast decrease their number by promoting apoptosis. More recently, we showed that this direct effect likely results from decreased expression and activity of mitochondria complex I genes, “oxidative phosphorylation” and respiration (oxygen consumption rate) and it requires Bak and Bax, two members of the Bcl-2 family of proteins that are critical for mitochondrial apoptotic death ^(3)^. In the cortical bone compartment, however, estrogens decrease osteoclast number indirectly by suppressing the expression of pro-osteoclastogenic factors produced by cells of the mesenchymal lineage. In support of this latter mechanism of action, we have also recently shown that in mice with conditional deletion of *Cxcl12* in *Prrx1* cells, the loss of cortical, but not trabecular, bone mass caused by estrogen deficiency is attenuated ^(4)^.

Proteolytic breakdown of extracellular matrix components plays an important role during bone remodeling. Collagen 1 and 2, the most abundant extracellular matrix components of bone and cartilage, are recycled via the activity of matrix metalloproteinase (MMP) family of enzymes. MMP-13 is highly expressed in terminally differentiated hypertrophic chondrocytes and osteoblasts. Also, a mutation of the human *Mmp-13* gene causes the Missouri variant of spondyloepimetaphyseal dysplasia (SEMD), a disorder characterized by abnormal development and growth of vertebrae and long bones ^(5)^. Studies of mice with global or conditional *Mmp-13* deletion have further elucidated the role of this metalloproteinase on the skeleton ^(6,7)^. MMP-13 deficiency in chondrocytes alters growth plate architecture and in osteoblasts/osteocytes increases trabecular bone mass ^(7,8)^. Several lines of evidence have implicated metalloproteinase in bone resorption. Indeed, the stimulation of bone resorption by parathyroid hormone (PTH) requires collagenase cleavage of type I collagen ^(9)^. In addition, MMP-13 stimulates osteoclast differentiation and activation in breast tumor bone metastases ^(10)^ and is involved in the osteolytic lesions of multiple myeloma ^(11)^. Interestingly, the action of MMP-13 in myeloma results from its ability to promote osteoclast fusion by up-regulating the fusogen DC-STAMP, independently of its enzymatic activity.

Notably, MMP-13 mRNA and protein increase in the osteoblasts of ovariectomized rats; and inhibition of metalloproteinase activity attenuates the loss of bone mass induced by estrogen deficiency in mice ^(12,13)^. This evidence has suggested that suppression of *Mmp-13* by estrogens may contribute to their bone protective effects. In the work presented here, we have generated mice lacking *Mmp-13* in mesenchymal lineage cells to functionally interrogate whether MMP-13 produced by these cells plays a role in the loss of bone mass caused by estrogen deficiency.

## Materials and Methods

### Animal Experimentation

Mice with conditional deletion of ERα using *Osx1-* and *Prrx1-Cre* and respective littermates were generated as previously described ^(14)^. Mice with conditional deletion of *Mmp-13* in the mesenchymal lineage were generated by a two-step breeding strategy. Hemizygous *Prrx1-cre* transgenic male mice (B6.Cg-Tg(*Prrx1-cre*)1Cjt/J; Jackson Laboratories, stock #5584) were crossed with female *Mmp-13* floxed (^f/f^) mice (FVB.129S-*Mmp-13^tm1Werb^*/J, Jackson Laboratories, stock # 005710) to generate mice heterozygous for the *Mmp-13* floxed allele with and without the *Cre* allele. These mice were intercrossed to generate *Mmp-13*^f/f^ (used as control) and *Mmp-13*^ΔPrrx1^ mice. Offspring were genotyped by PCR using the following primer sequences: TGA TGA CGT TCA AGG AAT TCA GTT T, wild-type, product size 572 bp, CCA CAC TGC TCG ACA TTG, mutant, product size 372 bp and GGT GGT ATG AAC AAG TTT TCT GAG C, heterozygote, product size 372 bp and 572 bp. Offspring from all genotypes were tail-clipped for DNA extraction at the time of weaning (21 days) and then group-housed with same sex littermates. Mice were maintained at a constant temperature of 23°C, a 12-hour light/dark cycle, and had access to food and water *ad libitum*. All mice used in these experiments were obtained from the same group of breeders in 2 consecutive breeding cycles separated by 30 days. *Mmp-13*^f/f^ and *Mmp-13*^ΔPrrx1^ littermate male mice were harvested and analyzed at 16 weeks (13 *Mmp-13*^f/f^ and 12 *Mmp-13*^ΔPrrx1^) and 24 weeks of age (14 *Mmp-13*^f/f^ and 10 *Mmp-13*^ΔPrrx1^). Twenty-week-old female *Mmp-13*^ΔPrrx1^ mice and *Mmp-13*^f/f^ littermates were either OVX or sham-operated after being stratified by body weight (fifteen *Mmp-13*^f/f^ and 15 *Mmp-13*^ΔPrrx1^ were sham operated; 12 *Mmp-13*^f/f^ and 12 *Mmp-13*^ΔPrrx1^ were OVX). Specifically, within each genotype, mice were sorted from low to high weight values. Mice were then assigned the numbers 1 and 2, successively. All animals with the same number were assigned to the same group. Weight means and standard deviation for each group were calculated and compared by *t*-test to assure that means were similar. Surgeries were performed in the morning under sedation with 2% isoflurane, as previously described ^(15)^. Mice were injected with calcein (Sigma-Aldrich, C0876; 35 mg/kg body weight) 7 and 3 days before euthanasia for quantification of bone formation. Animals were euthanized 8 weeks after surgery and tissues dissected for further analyses. Whole body weight was measured 2 days before surgery, before calcein injections and before euthanasia. Uterine weights were obtained to confirm depletion of sex steroids. Investigators were blinded during animal handling and endpoint measurements. No adverse events occurred during surgeries, calcein injections and harvest procedures. For bone strength measurements 13 *Mmp-13*^f/f^ and 14 *Mmp-13*^ΔPrrx1^ females with 26 weeks old were euthanized and femurs were harvested. The Institutional Animal Care and Use Committees of UAMS and the Central Arkansas Veterans Healthcare System approved the animal protocols used in these studies.

### Bone imaging

Right femurs and tibias from male and female *Mmp-13*^f/f^ and *Mmp-13*^ΔPrrx1^ mice were dissected, cleaned from adherent muscle and fixed in 10% Millonig’s formalin, dehydrated, and kept in 100% ethanol at 4°C. Two female tibiae were damaged during the harvest procedure and discarded. Femur and tibia lengths were measured with a micrometer followed by micro-CT analysis using a μCT40 (Scanco Medical, Bruttisellen, Switzerland). Bones were scanned at 12 μm nominal isotropic voxel size, 500 projection (medium resolution, E=55 kVp, I=72 μA, 4W, integration time 150 ms and threshold 200 mg/cm^3^), integrated into 3-D voxel images (1024*x*1024 pixel matrices for each individual planar stack) from the distal epiphysis of the femur and the proximal epiphysis of the tibia towards the mid-diaphysis to obtain a number of slices variable between 650 and 690. Cortical thickness, total and medullary area of the diaphysis were determined using 18 slices at the femur and tibia mid-shafts and cortical thickness of the distal metaphysis was determined analyzing slices 300 to 350 (counting from midshaft region). Cortical analysis was performed with a threshold of 200 mg/cm^3^. Two-dimensional evaluation of trabecular bone was performed on contours of the cross sectional acquired images excluding the primary spongiosa and cortex. Contours were drawn starting 8-10 slices away from the growth plate from the distal metaphysis towards the diaphysis of the femur to obtain 151 slices (12 μm/slice), or 8-10 slices away from the growth plate of the proximal metaphysis towards the diaphysis of the tibia to obtain 120 slices (12 μm/slice). For all trabecular bone measurements contours were drawn every 10 to 20 slices. Voxel counting was used for bone volume per tissue volume measurements and sphere filling distance transformation indices were used for trabecular microarchitecture with a threshold value of 220 mg/cm^3^, without pre-assumptions about the bone shape as a rod or plate. Micro-CT measurements were expressed in 3-D nomenclature as recommended by ASBMR standard guidelines ^(16)^.

### Histology and histomorphometry analysis

After μCT analysis, the right femurs from female *Mmp-13*^f/f^ and *Mmp-13*^ΔPrrx1^ mice were embedded undecalcified in methyl methacrylate. Calcein labels and osteoclasts were quantified on both endocortical surfaces in 5 μm thick longitudinal/sagittal sections using the OsteoMeasure Analysis System (OsteoMetrics, Inc. Atlanta, GA). Osteoclasts were stained for tartrate-resistant acid phosphatase (*Acp5*) using naphthol AS-MX and Fast Red TR salt (Sigma-Aldrich). The following primary measurements were made: bone surface (BS), single label surface (sL.S), double label surface (dL.S), inter-label thickness (Ir.L.Th), osteoclast number (N.Oc, /μm), and osteoclast surface (Oc.S, %). The following derived indices were calculated: mineralizing surface (MS, %), mineral apposition rate (MAR, μm/d), and bone formation rate (BFR, μm^3^/μm^2^/d). The referent for Oc.S, MS and BFR was BS, and for N.Oc B.Pm. During dynamic histomorphometric analysis we noticed that 5 mice were missing one of the calcein injections since no double labels were observed (2 mice in *Mmp-13*^f/f^ sham group, 1 mouse in the *Mmp-13*^ΔPrrx1^ sham group and 2 mice in *Mmp-13*^ΔPrrx1^ OVX group). These mice were excluded from the analysis. All histology measurements were made in a blinded fashion. We used the terminology recommended by the Histomorphometry Nomenclature Committee of the American Society for Bone and Mineral Research ^(17)^.

### Bone strength

Three-point bending tests were performed at room temperature, with the posterior femoral surface lying on lower supports at a 6.6 mm span. Load was applied to the anterior femoral surface by an actuator midway between the two supports, at a constant rate of 1 mm/min to failure (ElectroForce 5500, TA Instruments, New Castle, DE). Load (N) and displacement (mm) were recorded. The moment of inertia (MOI) in the midshaft of the femur was calculated using geometry measured from μCT scans (model μCT40, Scanco Medical). Yield stress, ultimate stress, and modulus of elasticity were calculated using the mechanical testing parameters, moment of inertia, and μCT measurements.

### Microarray Analysis

Raw signal intensity files from the BeadStudio of all samples – GFP-sorted Osx1^+^ cells without or with ERα, derived from calvaria cells of ERα^ΔGFP:Osx1^ mice or GFP:Osx1-Cre controls – were processed together by lumi package ^(18)^ under R suite software. Quantile normalization was performed to make data comparable across samples. Differential gene expressions between the two groups was evaluated by moderated Student’s t-test using limma package ^(19)^. The statistical P values were further adjusted for multiple testing using Benjamini-Hochberg method.

### Cell cultures

Bone marrow stromal cells from 6-month-old mice with conditional deletion of ERα using *Prrx1-Cre* and control littermates were obtained by flushing the tibias and femurs. Cells from 3 mice per group were pooled and cultured in α-MEM (Sigma) supplemented with 20% fetal bovine serum (FBS) (Sigma), 1% penicillin-streptomycin-glutamine (PSG) (Sigma) and 50 μg/mL ascorbic acid (Sigma) in 10 cm culture dishes for 5-7 days. Half of the medium was replaced every 3 days. Adherent bone marrow stromal cells were then re-plated in triplicate in 12 well plates at 2×10^5^ cells per well with ascorbic acid and 10 mM β‐glycerophosphate (Sigma) to perform qPCR assays. Calvaria cells from 3-4 day-old *Osx1-Cre* ERα deleted mice were isolated and cultured as described previously ^(14)^.

### RNA isolation and qPCR assay

The left femur and tibia shafts from female *Mmp-13*^f/f^ and *Mmp-13*^ΔPrrx1^ mice were flushed to remove the bone marrow, cleaned from adherent tissues and frozen in liquid N_2_. Frozen shafts were pulverized with a multi-well tissue pulverizer (BioSpec Products, Inc. Bartlesville, OK, USA) and frozen in Trizol at −80°C. Total RNA was isolated following the Trizol reagent method (Life Technologies, 15596). RNA from cultured cells from ERα *Osx1-* and *Prrx1-Cre* deleted mice and control littermates were extracted using the same methodology. RNA was reverse-transcribed using the High-Capacity cDNA Archive Kit (Applied Biosystems, Carlsbad, CA, USA). Taqman quantitative PCR was performed to determine mRNA levels of *Mmp-13 and ERa* using the Mm00439491_m1 and Mm00433148_mH primers respectively, manufactured by the TaqMan Gene Expression Assay service (Applied Biosystems). The mRNA expression levels were normalized to the house-keeping gene mitochondrial ribosomal protein S2 (*Mrsp2,* Mm00475528_m1) using the ΔCt method ^(20)^.

### Western blots

Bone marrow plasma from wild-type C57BL/6 mice was collected by removing both ends of femora and tibiae with a scalpel and then centrifuging the diaphyseal bone in a microcentrifuge tube at 5000 rpm for 30 s. The bone marrow cell pellet was then re-suspended in 90 μl of phosphate-buffered saline and then centrifuged at 8000 rpm for 1 min. The supernatant was transferred to a fresh tube and stored at −80°C until further analysis. Equal amounts of supernatant (40 μl) were incubated with SDS-PAGE sample buffer (125 mM Tris-HCl, pH 6.8, 4% SDS, 20% glycerol, 10% beta-mercaptoethanol, and 0.004% bromophenol blue) at 100°C for 5 min, loaded in each well, electrophoresed in 0.1 % SDS, X % gradient acrylamide gels, and transferred electrophoretically onto PVDF membranes (Millipore). The membranes were blocked in 5 % fat-free milk/Tris-buffered saline for 120 minutes and incubated with a mouse monoclonal antibody against MMP-13 (MilliporeSigma, MAB3321, 1:500), followed by a secondary antibody (Cell signaling technology) conjugated with horseradish peroxidase. The MMP-13 antibody specifically reacts with precursor and active forms of human or mouse MMP-13. Bound antibodies were detected with ECL reagents (Millipore) and imaged and quantified with a VersaDoc imaging system (Bio-Rad).

### Statistical analysis

For statistical analysis and preparation of graphs we used GraphPad Prism 9 software. Data are presented as dot plots with the central line representing the mean and error bars representing standard deviation. After determining that data were normally distributed and exhibited equivalent variances among groups, mean values were compared by 2-way ANOVA with Bonferroni multiple comparison test or by unpaired student t-test. When data were not normally distributed, Runk Sum test was used instead of Student’s t-test (Figs. 1C-D). In the case of ANOVA, data were *l*og10 transformed to achieve normal distribution and equivalence of variance and subjected to post hoc analysis of mean differences after Tukey adjustment (Figs. 2C, 4A, 4C-E, 5A-C, 5E) (). Outliers were identified and removed by the ROUTE method with a Q=1 % or by the Grubbs test with α = 5 % when data is normally distributed. Exact *p* values are shown for relevant comparisons. In line with the recommendations of the American Statistical Association, summarized by Amrhein et al ^(21)^, a threshold value of *p* was not used to define a statistically significant effect.

**Figure 1.**
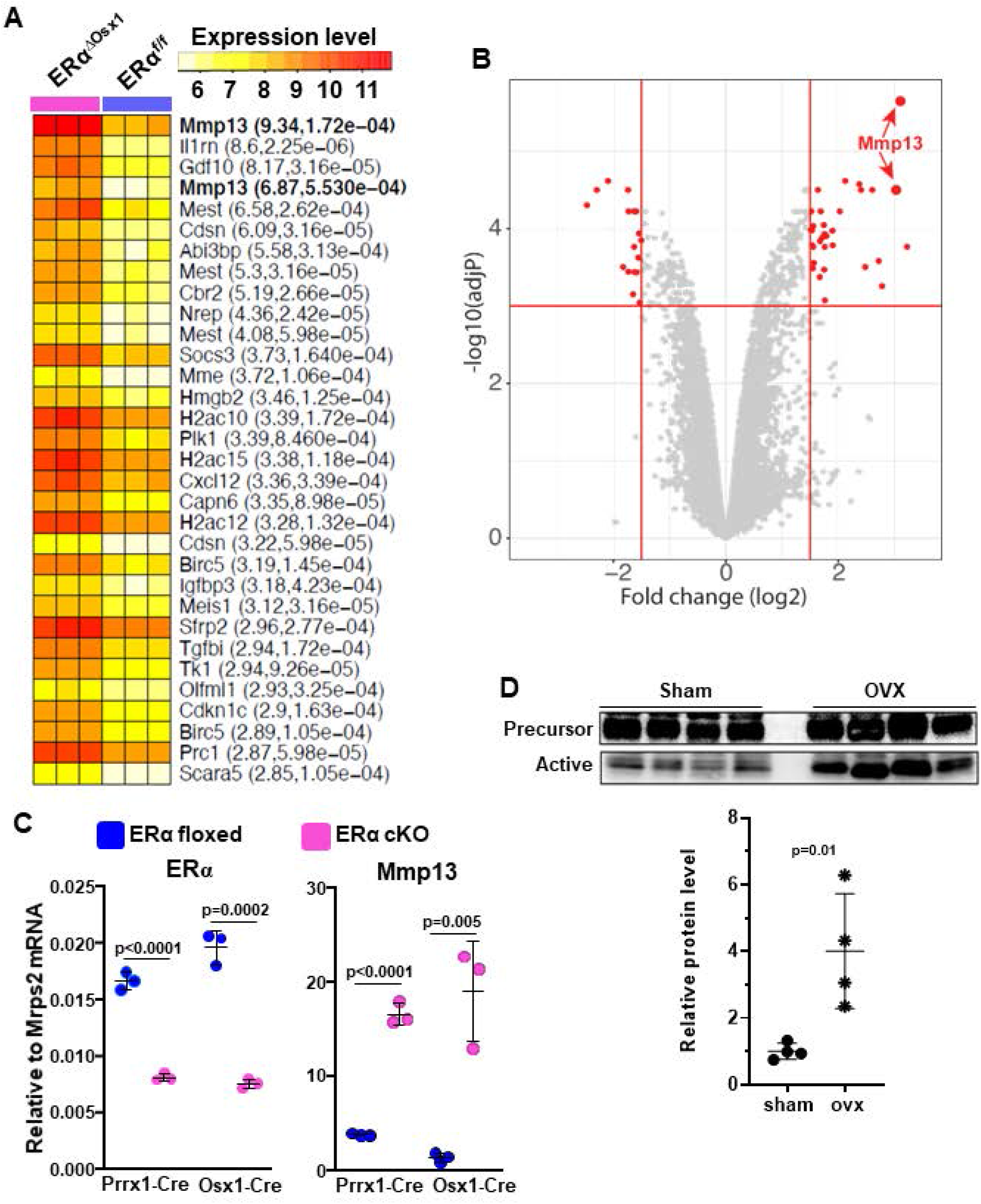
*Mmp-13* is downregulated by estrogens. (A) Microarray analysis of GFP-sorted Osx1^+^ cells without or with ERα, derived from calvaria cells of ERαΔOsx1 mice or Osx1-Cre controls. Heat map shows the normalized expression values of the top highly up-regulated genes (log2 fold change > 1.5 and adjust P values < 0.001) of individual sample. Gene names are shown on the right-side of the heatmap with fold change and adjusted p values. *Mmp-13* gene, which has 2 probe sets is in bold letters. (B) Volcano plots showing the profiles of differential gene expression. The gene that passed that cutoff (log2 fold change > 1.5 and adjust P values < 0.001) are represented by red color dots. *Mmp-13* gene, which has 2 probe sets is pointed to by the red arrows. (C) Relative mRNA levels of ERα and *Mmp-13* derived from *Prrx1-Cre* bone marrow stromal cells and *Osx1-Cre* calvaria cells from ERα deleted mice (ERα^ΔPrrx1^ and ERα^ΔOsx1^, respectively) and littermate controls (ERα^f/f^). (D) Protein levels of the precursor and active form of MMP-13 protein on bone marrow plasma of ovariectomized and sham operated wild type mice. The Western blots depict both precursor (inactive form) and active form from the same loaded sample (each sample is a pool of 3-4 animals from either group); the ratio of the latter over the former is shown in the dot plot. Data represent mean ± S.D. (n=14-12 mice/group); p values by student t-test.

**Figure 2.**
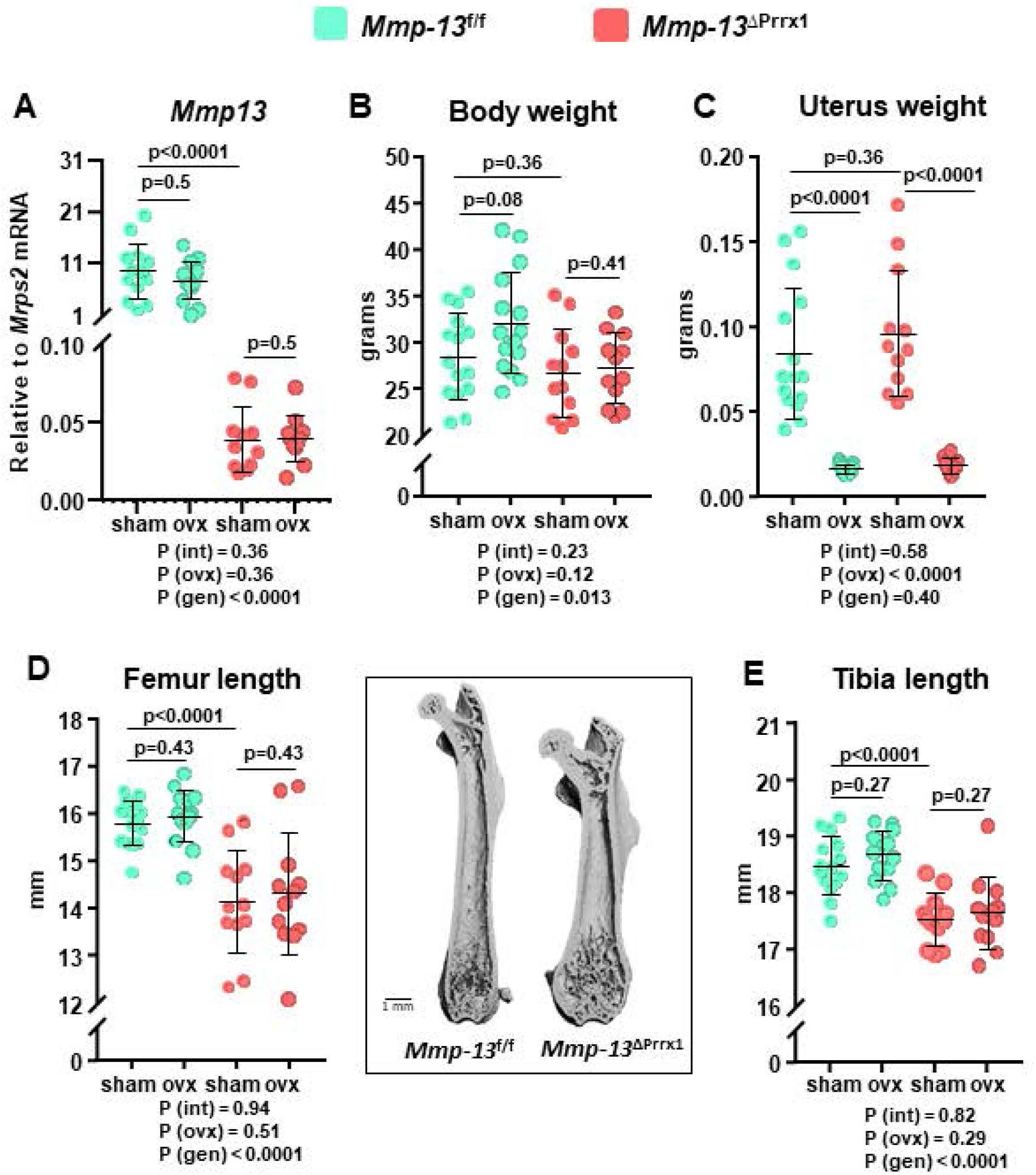
*Mmp-13* deletion in mesenchymal progenitors decreases the lengths of both femur and tibia. Female mice with *Mmp-13* deletion in *Prrx1* expressing cells (*Mmp-13*^ΔPrrx1^) and control littermates (*Mmp-13*^f/f^) where either ovariectomized (OVX) or sham-operated (Sham) at 5 months of age and euthanized 8 weeks later. (A) *Mmp-13* relative mRNA expression in femur shafts devoid of bone marrow by qRT-PCR. (B) Body weight and (C) uterine weight measured at euthanasia. (D) Quantification of femur length and representative micro-CT images of longitudinal femur sections of *Mmp-13*^f/f^ and *Mmp-13*^ΔPrrx1^ mice and (E) quantification of tibia length. Data represent mean ± S.D. (n=12-15 mice/group); P values by 2-way ANOVA. Interaction terms generated by 2-way ANOVA analysis are shown below each graph (P value of the interaction, P value of the ovariectomy and P value of the genotype).

## RESULTS

### *Mmp-13* expression is downregulated by ERα

To identify estrogen target genes that regulate osteoclastogenesis indirectly, i.e., via actions on cells of the mesenchymal lineage, we performed microarray analysis of GFP sorted Osx1+ cells without or with ERα, derived from the calvaria cells of ERα^f/f^;GFP:*Osx1-Cre* mice or GFP:*Osx1-Cre* controls. The highest up-regulated gene in ERα deleted mesenchymal/stromal cells encodes the matrix metalloproteinase 13 (*Mmp-13*), as shown in the heat map (Figure 1A) and in the volcano plot (Figure 1B). The microarray findings from the GFP sorted ERα deleted *Osx1+* calvaria cells were confirmed by qPCR and reproduced in cultures of ERα deleted *Prrx1+* bone marrow stromal cells derived from ERα^f/f^;*Prrx1-Cre* mice (Figure 1C). Moreover, the active form of MMP-13 protein levels was 4-fold higher in the BM plasma of OVX C57BL/6 mice, as compared to sham controls (Figure 1D). In line with our findings, ERα regulates the *Mmp-13* promoter activity in synoviocytes ^(22)^.

### *Mmp-13* deletion in mesenchymal progenitors decrease the length of the femur and tibia

To elucidate the role of *Mmp-13* in bone homeostasis *in vivo*, we next generated mice with conditional deletion of *Mmp-13* in mesenchymal progenitors expressing *Prrx1 (Mmp-13*^ΔPrrx1^) and used floxed mice (*Mmp-13*^f/f^) as control. The *Prrx1-cre* transgene targets early limb bud and a subset of craniofacial mesenchymal stem cells. We did not detect a skull phenotype. All measurements were made in the femur and/or tibia. Please note that in the following description of the results, the p values from two-way ANOVA analysis are provided below each graph. The expression of the *Mmp-13* mRNA in femur and tibia shafts was dramatically decreased in the *Mmp-13*^ΔPrrx1^ mice (Figure 2A), establishing the effectiveness of the deletion. Body weight and uterine weight were not affected by the *Mmp-13* deletion (Figures 2B and 2C). However, femur and tibia length was decreased (Figures 2D and 2E).

In contrast to the microarray data of figure 1, we did not detect a change of the mRNA levels of *Mmp-13* in the OVX *Mmp-13*^f/f^ or *Mmp-13*^ΔPrrx1^ mice in the osteocyte-enriched bone marrow-devoid preparations we used for this measurement (Figure 2A). As expected, OVX increased body weight in *Mmp-13*^f/f^ mice (Figure 2B) and decreased uterine weight in *Mmp-13*^f/f^ and *Mmp-13*^ΔPrrx1^ mice (Figures 2C). Femur and tibia length was not affected by OVX in either genotype (Figures 2D and 2E).

### *Mmp-13* deletion increases cortical bone and attenuates the cortical bone loss caused by OVX

*Mmp-13* deletion in *Prrx1* cells increased cortical thickness and cortical area in the femoral diaphysis (Figure 3A and 3B). This effect was due to a decrease in the medullary area (Figure 3C) while total area did not change (Figure 3D). The increase in cortical thickness with *Mmp-13* deletion was less marked in the distal metaphysis of the femur (Figure 3E) and in the diaphysis of the tibia (Figure 3F).

**Figure 3.**
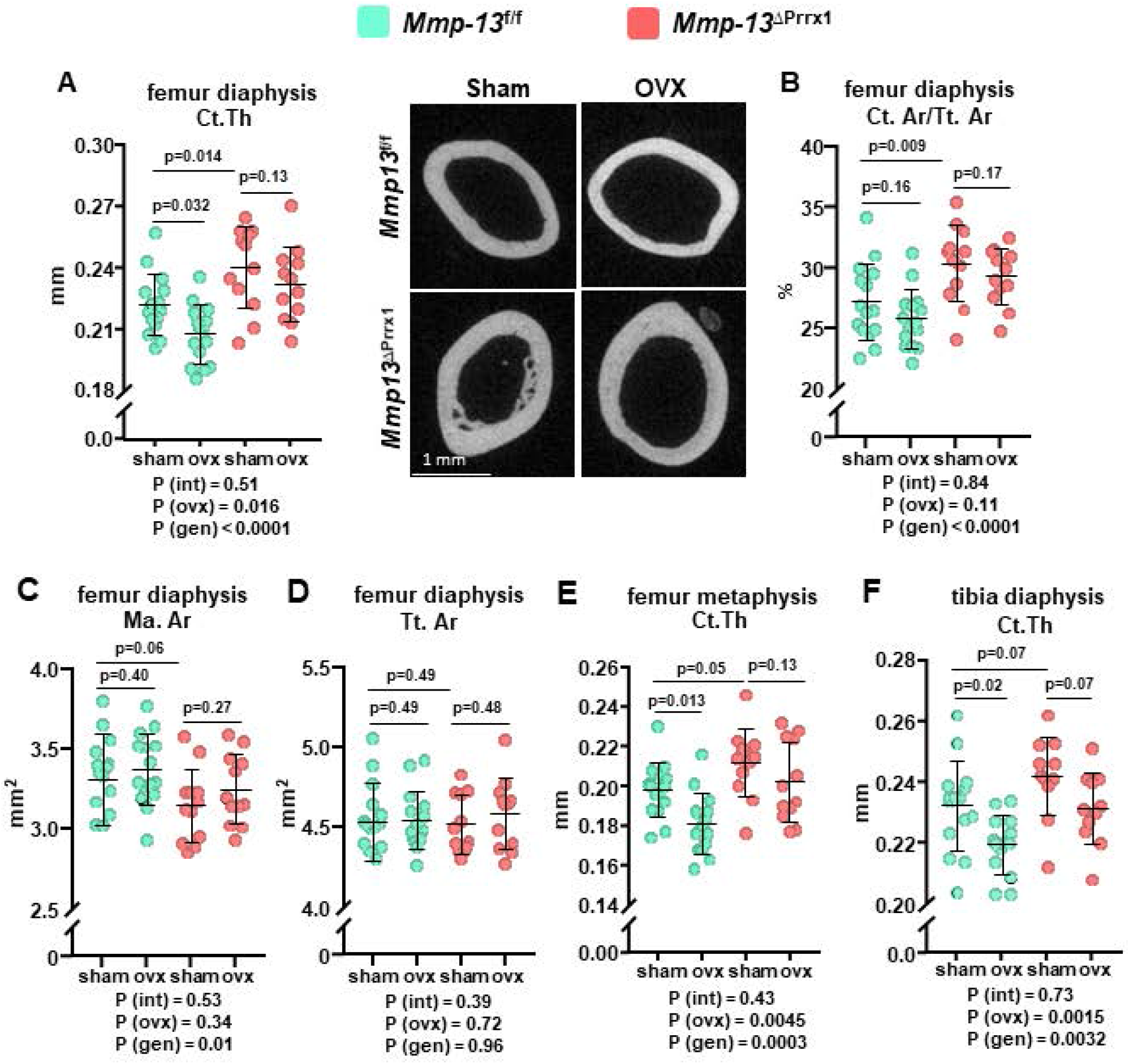
*Mmp-13* deletion increases cortical bone and attenuates the cortical bone loss caused by ovariectomy. Femur and tibia cortical bone were evaluated by micro-CT. (A) Cortical thickness measured in diaphysis (mid-shaft) and representative micro-CT images of the same region of the femur. (B) Cortical area under total area, (C) medullary area and (D) total area measured in mid-shaft femur. (E) Cortical thickness measured at distal metaphysis of the femur. (F) Cortical thickness measured in the diaphysis (mid-shaft) of the tibia. Data represent mean ± S.D. (n=12-15 mice/group); P values by 2-way ANOVA. Interaction terms generated by 2-way ANOVA analysis are shown below each graph (P value of the interaction, P value of the ovariectomy and P value of the genotype).

OVX of the *Mmp-13*^f/f^ control mice caused a decrease in cortical thickness at the femoral diaphysis and distal metaphysis as well as the tibia diaphysis (Figure 3A, 3E and 3F). Consistent with our working hypothesis, the effects of OVX on cortical bone at the diaphysis and distal metaphysis of the femur and tibia diaphysis were attenuated in the *Mmp-13*^ΔPrrx1^ mice (Figures 3A, 3E and 3F). Together, these data suggest that *Mmp-13* deletion increases cortical bone mass in femur and tibia and prevents or attenuates the loss of cortical bone caused by estrogen deficiency.

### *Mmp-13* deletion increases trabecular bone but does not affect the loss of bone caused by OVX in this compartment

*Mmp-13* deletion in *Prrx1* cells increased trabecular bone volume in both the distal femur and proximal tibia by approximately 3.7- and 3-fold, respectively (Figure 4A and 4E). This increase was due to an increase in trabecular number (Figure 4B) and thickness (Figure 4C); and was mirrored by a decrease in trabecular separation (Figure 4D).

**Figure 4.**
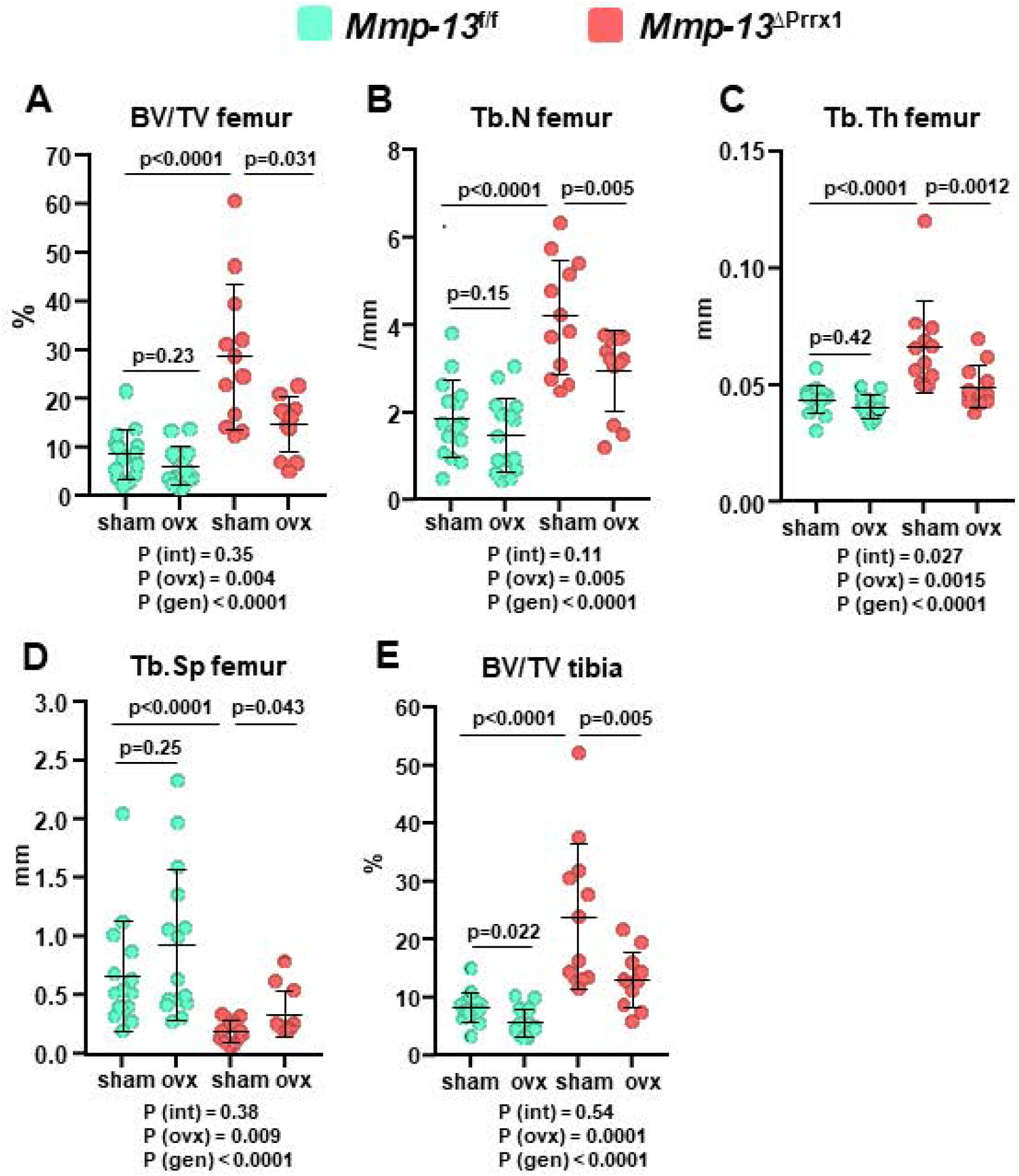
*Mmp-13* deletion increases trabecular bone but does not affect the loss of bone caused by ovariectomy in this compartment. Femur and tibia trabecular bone were evaluated by micro-CT. (A) Trabecular bone volume and (B-D) microarchitecture at the distal metaphysis of the femur. (E) Trabecular bone volume at the proximal metaphysis of the tibia. Data represent mean ± S.D. (n=12-15 mice/group); P values by 2-way ANOVA. Interaction terms generated by 2-way ANOVA analysis are shown below each graph (P value of the interaction, P value of the ovariectomy and P value of the genotype).

As seen before ^(23,24)^, at six-months of age estrogen sufficient female mice have very little trabecular bone mass remaining at the distal femur (Figure 4A). There was no discernible effect of the OVX at this site in *Mmp-13*^f/f^ mice (Figure 4A-D). However, we observed a loss of trabecular bone mass in both the femur and tibia of the OVX *Mmp-13*^ΔPrrx1^ mice (Figure 4A and 4E). Collectively, these data indicate that *Mmp-13* deletion increases trabecular bone mass but does not prevent the loss of trabecular bone caused by estrogen deficiency.

### *Mmp-13* deletion decreases osteoclast number in cortical bone

To elucidate the cellular mechanism by which the *Mmp-13* deletion increased cortical bone volume, we performed histomorphometric analysis of the endocortical surface of the femur. *Mmp-13* deletion caused an approximately 50% reduction in osteoclast number and surface (Figure 5A-C), but had no effect on mineral apposition rate (MAR), mineralized surfaces (MS) or bone formation rate (BFR) (Figure 5 D-G). These findings suggest that a decrease of osteoclast number and thereby resorption are responsible for the increase of cortical bone.

**Figure 5.**
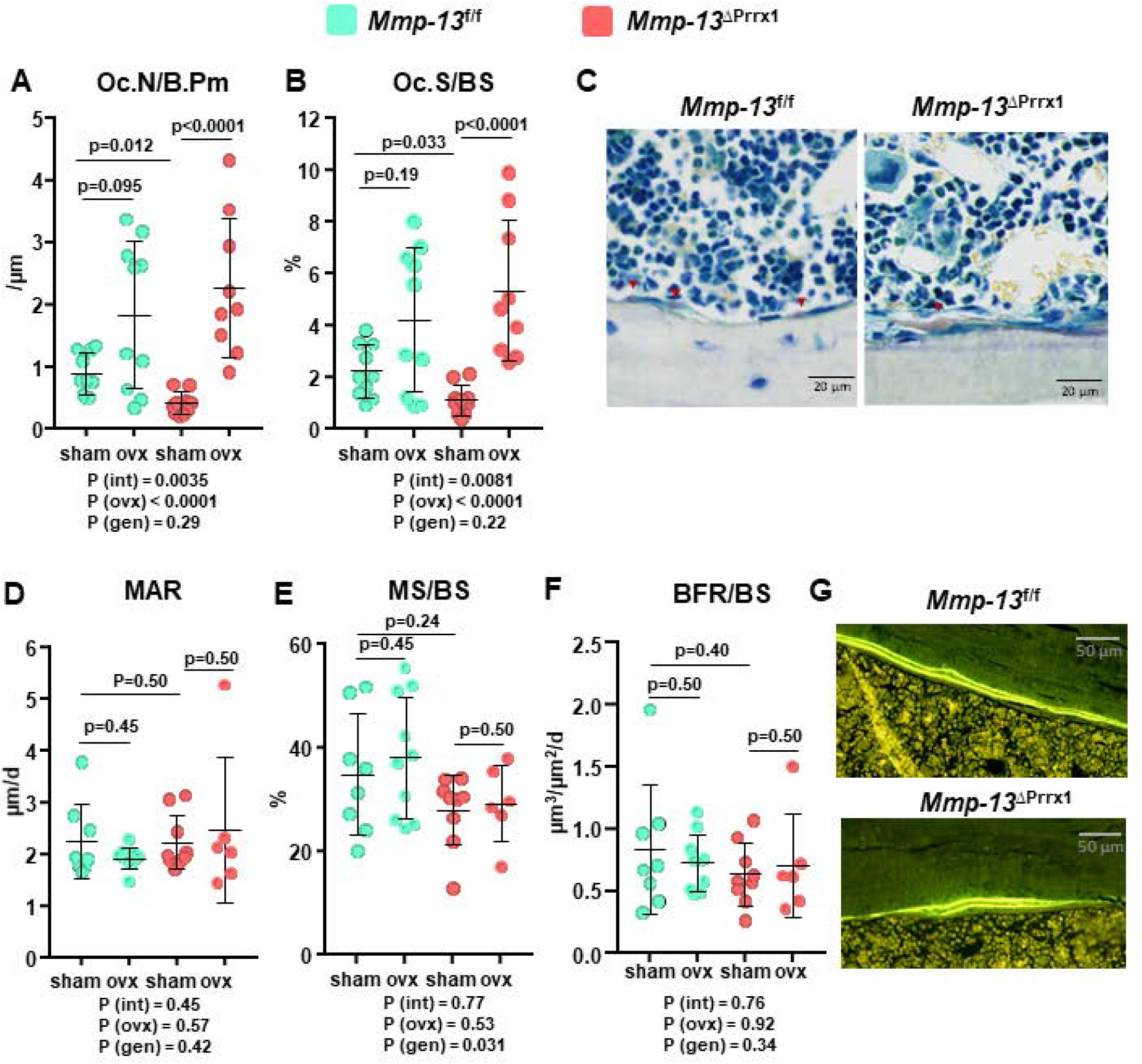
*Mmp-13* deletion decreases osteoclast number at the endocortical surface of the femur. Histology and dynamic histomorphometry were evaluated at the endocortical bone surface in longitudinal undecalcified femur sections from 6-month-old female mice sham or OVX operated. (A) Osteoclast number per bone perimeter, (B) osteoclast surface per bone surface, and (C) representative microphotographs of osteoclasts in sections from sham animals stained with tartrate-resistant acid phosphate (*Acp5*). (D) Mineral apposition rate, (E) mineralizing surface and (F) bone formation rate, and (G) representative photomicrographs of cortical bone labeled with calcein in sections from sham animals. Data represent mean ± S.D. (n=6-10 mice/group): P values by 2-way ANOVA. Interaction terms generated by 2-way ANOVA analysis are shown below each graph (p value of the interaction, p value of the ovariectomy and p value of the genotype).

OVX of both *Mmp-13*^f/f^ and *Mmp-13*^ΔPrrx1^ mice resulted in the expected increase in osteoclast number and surface (Figure 5A-B), while MAR, MS and BFR were not affected (Figure 5D-F). Surprisingly, in the OVX *Mmp-13*^ΔPrrx1^ mice the increase in osteoclast number and surface was greater (5.5-fold) as compared to the OVX-induced increase in the *Mmp-13*^f/f^ mice (2-fold).

### *Mmp-13* deletion increases whole-bone strength of the femur

It has been previously reported that *Mmp-13*−/− mice have increased cortical bone fragility ^(25)^. To examine bone strength in female *Mmp-13*^ΔPrrx1^ mice we performed three-point bending of the femur (Figure 6A). Despite the thicker cortices in female *Mmp-13*^ΔPrrx1^ mice, the moment of inertia was not different from the controls (Figure 6B). Nonetheless, the yield load, peak load, and stiffness were all higher in *Mmp-13*^ΔPrrx1^ mice (Figure 6C). With respect to the estimated material properties, female *Mmp-13*^ΔPrrx1^ mice had increased yield stress and ultimate stress but the same modulus as compared to control mice (Figure 6D). Material density determined by micro-CT was also not different (Figure 6E). Therefore, and in contrast to a previous report, deletion of *Mmp-13* led to an increase in bone structural and material properties.

**Figure 6.**
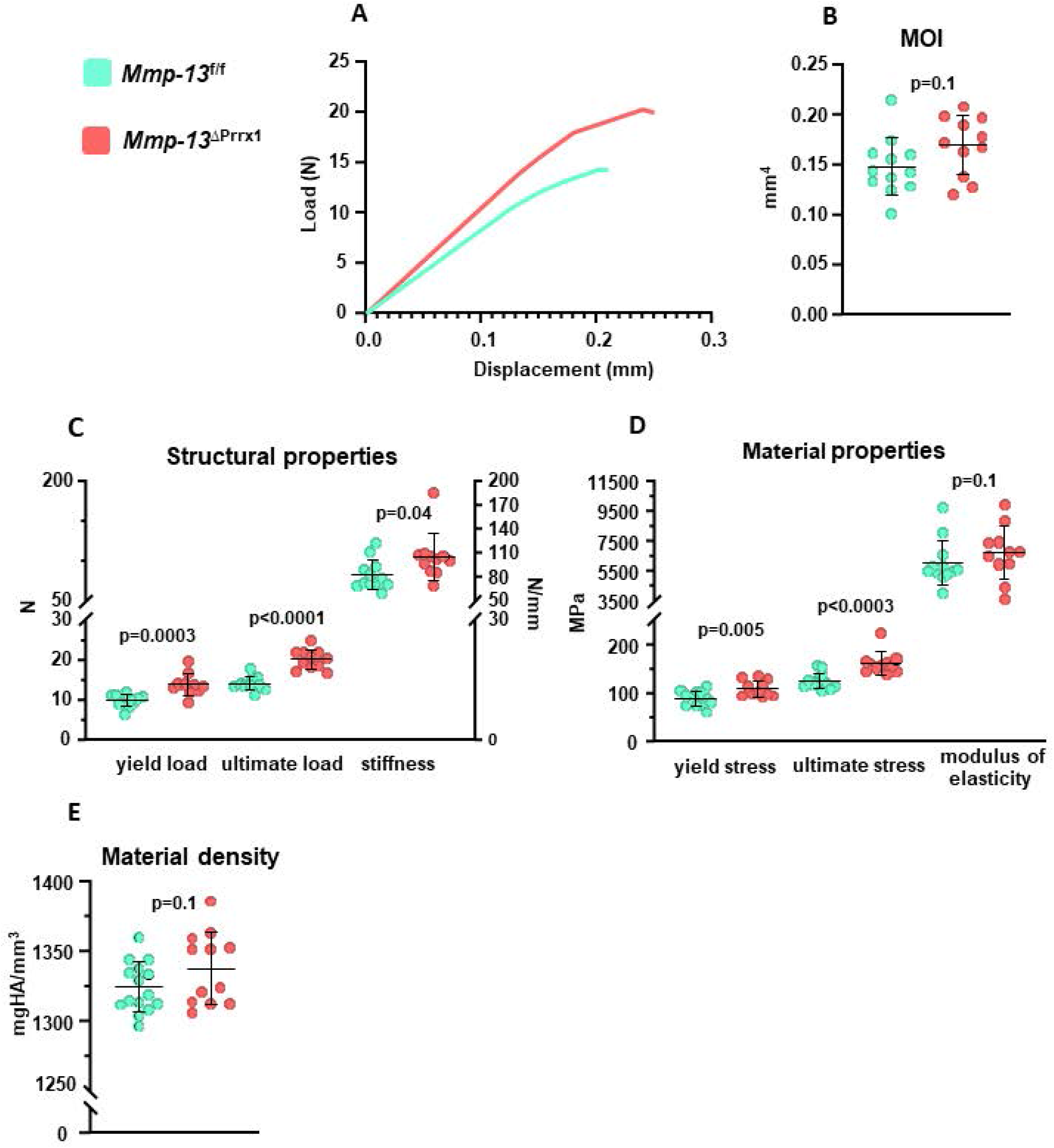
*Mmp-13* deletion increase whole-bone strength in female mice. Femurs from 6-month-old *Mmp-13*^f/f^ and *Mmp-13*^ΔPrrx1^ female mice were tested for bone strength by 3-point bending. (A) Representative load *versus* displacement curve showing that femur of *Mmp-13*^ΔPrrx1^ mice are more resistant than control littermates. (B) Moment of inertia, (C) Structural properties, and (D) Material properties. (E) Material density by micro-CT. Data represent mean ± S.D. (n=12-11 mice/group); P values by student t-test.

### Similar to females, *Mmp-13* deletion in males increases trabecular bone mass and whole-bone strength of the femur, but has no effect on cortical bone

The bone phenotype of sex steroid sufficient male *Mmp-13*^ΔPrrx1^ and *Mmp-13*^f/f^ mice was analyzed at 4 and 6 month of age. Body weight was not affected by the *Mmp-13* deletion at either age (Figure 7A), but femur length decreased at 4 and 6 months of age (Figure 7B), as it did in females. In difference to females, cortical thickness in male mice was not affected by the *Mmp-13* deletion (Figure 7C). Trabecular bone volume increased 1.4- and 1.6-fold at 4 and 6 months, respectively (Figure 7D), but this increase was lower compared to the one we observed in *Mmp-13*^ΔPrrx1^ female mice. The increased trabecular bone volume was associated with an increase in the number of trabeculae and a decrease in trabecular separation (Figure 7E and G), but no change in trabecular thickness (Figure 7F).

**Figure 7.**
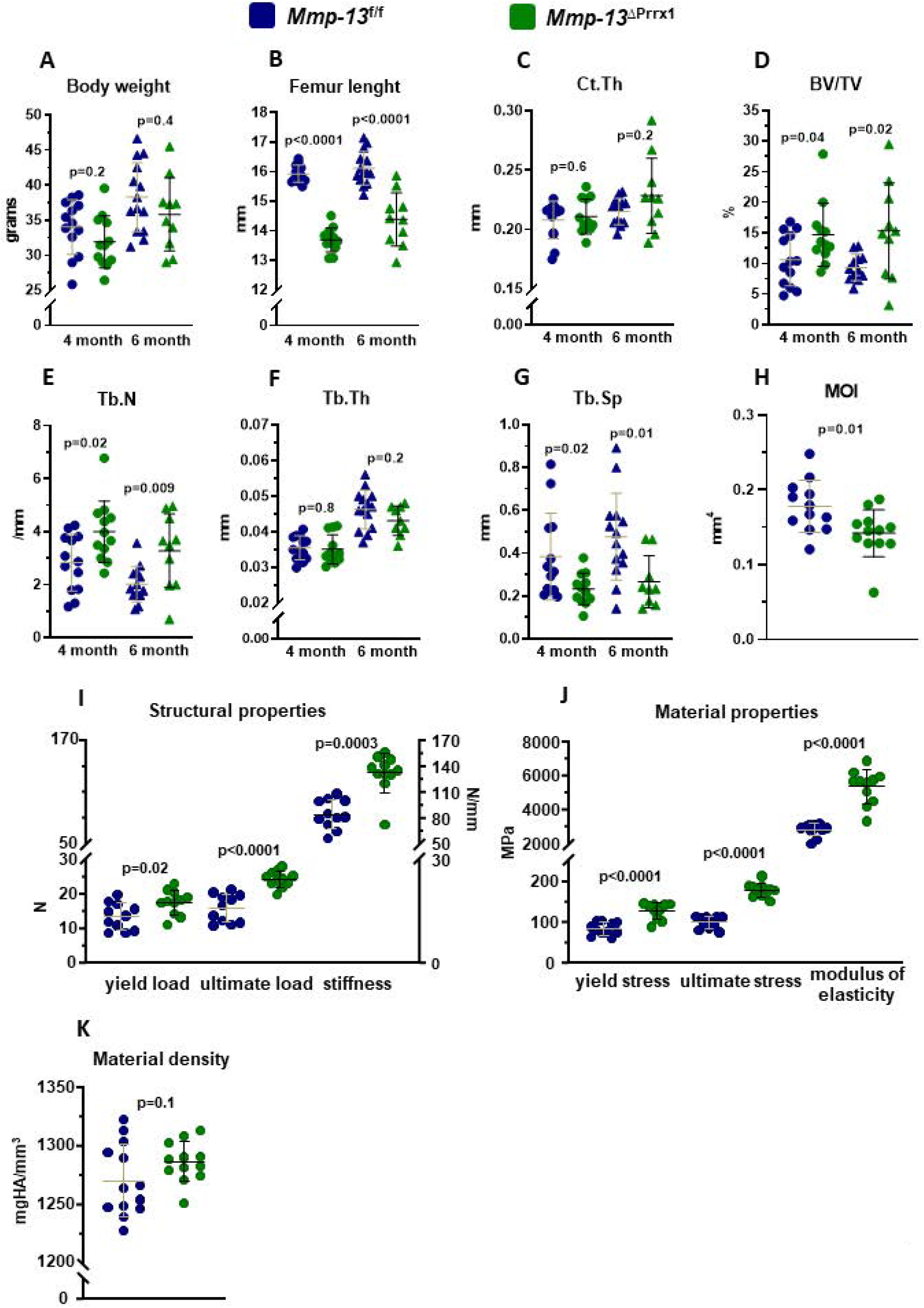
*Mmp-13* deletion in males increases trabecular bone mass and whole-bone strength. Male mice with *Mmp-13* deletion in Prrx1 expressing cells (*Mmp-13*^ΔPrrx1^) and control littermates (*Mmp-13*^f/f^) where euthanized at 4 and 6 month of age. (A) Whole body weight and (B) femur length measured with a micrometer. (C) Cortical thickness at mid-shaft femur. (D) Trabecular bone volume and (E-G) microarchitecture at the distal metaphysis of the femur by micro-CT. (H-J) Three-point bending test in femurs from 4-month-old *Mmp-13*^f/f^ and *Mmp-13*^ΔPrrx1^ male mice. (H) Moment of inertia, (I) structural properties of the femurs, including yield load, ultimate load and, stiffness, (J) material properties of the femurs, including yield stress, ultimate stress and modulus of elasticity. (K) Material density by micro-CT. Data represent mean ± S.D. (n=14-10 mice/group); P values by student t-test.

Finally, similar to females, three-point bending of the femur in male *Mmp-13*^ΔPrrx1^ mice revealed higher bone strength including an increase in stiffness and modulus (Figure 7H-J), with no change in material density (Figure 7K); but unlike females the moment of inertia was decreased in males (Figure 7H).

## DISCUSSION

In this paper we show that the highest upregulated mRNA in mesenchymal lineage cells lacking ERα encoded MMP-13. In estrogen sufficient (sham-operated) *Mmp-13*^ΔPrrx1^ mice cortical thickness and trabecular bone volume in the femur and tibia were increased as compared to littermate controls, while femur and tibia length was decreased. These bone phenotypic changes were associated with a decrease in osteoclast number, but no changes in bone formation. Moreover, the loss of cortical bone caused by OVX in the femur and tibia was attenuated in the *Mmp-13*^ΔPrrx1^ mice. The effect of OVX on trabecular bone, on the other hand, was not affected. These results elucidate an important role of mesenchymal cell–derived MMP-13 on osteoclast number, bone resorption, and bone mass. We had previously shown that the OVX-induced loss of cortical, but not trabecular bone, was attenuated in mice with conditional *Cxcl*12 deletion in *Prrx1* expressing cells. Taken together, the functional genetic evidence obtained by these two studies suggests that increased production of mesenchymal cell-derived factors, such as CXCL12 and MMP-13, are important mediators of the adverse effect of estrogen deficiency on cortical, but not trabecular, bone.

The expression of MMP-13 was increased in calvaria or bone marrow derived cells from ERα conditional KO mice. In contrast, loss of estrogen with OVX did not alter the expression of MMP-13 in osteocyte-enriched cortical bone from the femur. These findings suggest that estrogens attenuate the expression of MMP-13 in stromal cells or osteoblasts, but not osteocytes. Others have shown before that the MMP-13 content of osteoblastic cells in the proximal tibia increases with OVX in the rat ^(12)^. Furthermore, in human articular chondrocytes, estradiol suppresses the expression of MMP-13 ^(26)^; and an increase in MMP-13 has been associated with the deleterious effects of estrogen deficiency in osteoarthritis and intravertebral disc degeneration ^(27)^. In contrast, estrogens may promote temporomandibular joint disorders via induction of MMP-9 and MMP-13 in fibrochondrocytes ^(28)^. In synoviocytes, ERα may regulate the expression of MMP-13 through the AP-1 transcriptional regulatory site ^(29)^. However, other transcription factor binding sites such as Runx, PEA-3 and p53 in conjunction with AP-1 appear to be critical for the transcriptional regulation of *Mmp-13* in other cells ^(30,31)^, perhaps explaining the different responses of this gene to estrogens in different bone cell types.

It has been suggested before, that stimulation of collagenase activity, particularly by MMP-13, acts as a coupling factor for the activation of osteoclasts ^(9)^. However, other reports suggest that MMP-13 can stimulate osteoclast activity independent of its enzymatic activity ^(11)^. *Mmp-13* null mice are resistant to the loss of bone mass caused by multiple myeloma, though the number of osteoclasts on bone was unaffected by the MMP-13 deletion ^(32)^. In co-cultures, stromal cells derived from *Mmp-13* null mice increase the number of osteoclast, however these osteoclasts are smaller and resorb less bone. This evidence along with our findings that ablation of *Mmp-13* in *Prrx1* expressing cells does not prevent the increase in osteoclast caused by OVX, supports the idea that in pathologic conditions MMP-13 may promote the bone resorbing activity of osteoclasts, not osteoclastogenesis.

In line with previous evidence from *Mmp-13* null mice ^(6)^, we found that *Mmp-13* deletion in the *Prrx1* targeted mesenchymal progenitors decreased the length of the femur and tibia in both male and female mice. This effect most likely results from the expansion of hypertrophic cartilage in the growth plate that occurs during development and growth and it is caused by the deletion of *Mmp-13* in growth plate chondrocytes ^(7)^, which are targeted by *Prrx1*-Cre. We also confirmed herein that *Mmp-13* deletion increases trabecular bone volume in the femur and tibia. This effect was seen in both sexes. However, *Mmp-13*^ΔPrrx1^ females exhibited a bigger increase than males. Deletion of *Mmp-13* in *Col1*-Cre or *Dmp1*-Cre targeted cells also increases trabecular bone mass, indicating that osteoblasts and/or osteocytes are the major source of MMP-13 responsible for this effect ^(7,8)^. Nonetheless, we cannot exclude the possibility that MMP-13 in cells of the mesenchymal lineage other than osteoblasts and osteocytes contributes to the changes in bone mass.

In the present report we show for the first time that *Mmp-13* deletion in females increases cortical thickness; and this effect is associated with a decrease in osteoclast number in the endocortical surface with no changes in bone formation. Interestingly, and similar to previous reports in male *Mmp-13* KO mice ^(25)^, cortical thickness was unaffected in our male *Mmp-13*^ΔPrrx1^ mice. Despite the increase in trabecular bone mass, we found no changes in osteoclast number or bone formation in trabecular bone in 6-month-old females. Nonetheless, Nakatani et al ^(33)^ have shown that the number of osteoclasts is severely decreased in the trabecular bone of 8-day-old *Mmp-13* KO mice. Thus, it is possible that osteoclast number was decreased at an earlier age in our mice. Collectively, this evidence indicates that MMP-13 is a potent stimulant of bone resorption, particularly in female mice.

It has been reported previously that male mice with global *Mmp-13* deletion or osteocyte-specific *Mmp-13* ablation have defective osteocyte perilacunar remodeling and decreased bone toughness ^(25)^. Albeit, the mice with the osteocyte-specific *Mmp-13* ablation exhibited incongruent changes in cortical bone biomechanical properties: decreased ultimate load but increased yield load and yield stress. The *Prrx1*-Cre transgene in our *Mmp-13*^ΔPrrx1^ mice has inexorably deleted *Mmp-13* in osteocytes. Yet, in contrast to these previous findings, both male and female *Mmp-13*^ΔPrrx1^ mice exhibited increased femoral bone strength. We have not performed an examination of the osteocyte canalicular network in our mice, but the increase in femoral strength we found in *Mmp-13*^ΔPrrx1^ mice argues against a biomechanically consequential change of the lacuna-canalicular network.

Cellular senescence is a hallmark of aging ^(34–37)^ and a state in which cells secrete an array of pro-inflammatory cytokines, chemokines and proteases, known collectively as the Senescence Associated Secretory Phenotype (SASP) ^(38,39)^. Work by us and others has shown that osteoprogenitors and osteocytes from old mice exhibit markers of senescence and SASP, including higher levels of MMP-13 and CXCL12 and that the adverse effects of aging on murine cortical bone are due, at least in part, to cellular senescence ^(40–42)^. As shown herein and in our previously published studies with *Cxcl12*^ΔPrrx1^ mice, both MMP-13 and CXCL12 contribute to the loss of cortical bone caused by estrogen deficiency. We find it intriguing that some of the same cytokines may be responsible for the increase in osteoclast number and loss of cortical bone caused by both estrogen deficiency and old age.

In conclusion, the work described herein, adds to and fully supports a long line of evidence that MMP-13 plays an important role, not only in bone development, but also during bone remodeling in postnatal life. These effects are evidently mediated by changes in osteoclast number and perhaps activity during the resorption phase of remodeling. Furthermore, *Mmp-13* is a target gene of ERα signaling in mesenchymal progenitors and their descendants, such as bone marrow stromal cells, osteoblast progenitors, and matrix synthesizing mature osteoblasts. Loss of the restraining effect of estrogens on MMP-13 in estrogen deficient states, such as OVX in mice and menopause in women, is therefore an important culprit of the increased resorption associated with these states. Importantly, even though mesenchymal cell–derived MMP-13 influences trabecular bone mass, it plays no role in the loss of trabecular bone caused by estrogen deficiency, highlighting the striking divergence of the cellular targets and downstream mediators of the effects of ERα activation by estrogens in cortical versus trabecular bone. Whether MMP-13 plays also a role in the SASP-mediated loss of cortical bone in old age needs to be functionally investigated in future studies.

## ACKNOWLEDGEMENTS

This work was supported by the Biomedical Laboratory Research and Development Service of the Veteran’s Administration Office of Research and Development [I01 BX001405 (SCM)], the NIH [R01 AR056679 (MA)] and [P20 GM125503], and the University of Arkansas for Medical Sciences Tobacco Funds and Translational Research Institute (1UL1RR029884). We thank S Berryhill and J Crawford for technical assistance and Katie Poe for help with the preparation of the manuscript.

## AUTHOR CONTRIBUTIONS

SCM and MA designed the experiments and FP, HNK, SI, LH, AW, and EM performed the experiments. FP, HGA, IN, MA and SCM analyzed the data and wrote the manuscript.

